# Prewhitening and Normalization Help Detect a Strong Cross-Correlation Between Daily Wastewater SARS-CoV-2 RNA Abundance and COVID-19 Cases in a Community

**DOI:** 10.1101/2022.12.16.520829

**Authors:** Min Ki Jeon, Bo Li, Doris Yoong Wen Di, Tao Yan

**Author notes:** Corresponding author: Tao Yan, University of Hawaii at Manoa, Department of Civil and Environmental Engineering, 2540 Dole Street, 383 Holmes Hall, Honolulu, HI 96822.

## Abstract

Wastewater surveillance is a promising technology for real-time tracking and even early detection of COVID-19 infections in communities. Although correlation analysis between wastewater surveillance data and the daily clinical COVID-19 case numbers has been frequently conducted, the importance of stationarity of the time-series data has not been well addressed. In this study, we demonstrated that strong yet spurious correlation could arise from non-stationary time-series data in wastewater surveillance, and data prewhitening to remove trends helped to reveal distinct cross-correlation patterns between daily clinical case numbers and daily wastewater SARS-CoV-2 concentration during a lockdown period in 2020 in Honolulu, Hawaii. Normalization of wastewater SARS-CoV-2 concentration by the endogenous fecal viral markers in the same samples significantly improved the cross-correlation, and the best correlation was detected at a two-day lag of the daily clinical case numbers. The detection of a significant correlation between daily wastewater SARS-CoV-2 RNA abundance and clinical case numbers also suggests that disease burden fluctuation in the community should not be excluded as a contributor to the often observed weekly cyclic patterns of clinical cases.

**Water impact:** Wastewater surveillance represents an emerging water technology with significant human health benefits. The study demonstrated that non-stationary time-series data could lead to spurious correlation, highlighting the need for prewhitening. Normalization strategies could alleviate variations in sample collection and analyses, which is useful for detecting actual underlying relationships between wastewater surveillance data and clinical data.

## 1. Introduction

Since the outbreak of COVID-19 pandemic in late 2019 ^1^, wastewater surveillance has been explored as a new way to monitor the spread of SARS-CoV-2 in human communities. Many studies have shown the presence of SARS-CoV-2 viral particles or genomic RNA in bodily wastes, including feces ^2^, urine ^3^, and respiratory fluids ^4, 5^, in both symptomatic and asymptotic patients. In particular, asymptomatic infections are now known to account for a large percentage of total COVID-19 infections ^6, 7^, and also shed SARS-CoV-2 virus in feces ^8, 9^. Since wastewater collects human wastes from all individuals in the wastewater service area and hence can provide comprehensive information on COVID-19 infection in the community. This enables a unique advantage of wastewater surveillance in that it can potentially capture the “actual” infection rates, including the asymptomatic or mildly symptomatic patients in the community who are less likely to seek clinical testing.

Wastewater surveillance may also be able to provide real-time tracking and even early detection of infectious in the communities. The sources of SARS-CoV-2 viral shedding to wastewater include mainly feces and partially saliva and sputum due to their high shedding probability and the possibility of entering the sewers ^10^. It is known that COVID-19-infected patients start to shed SARS-CoV-2 virus in feces during the incubation period between the infection and the symptom onset ^11^ and the peak of SARS-CoV-2 viral concentration is reported generally at the beginning of the symptom onset ^12, 13^. In addition, model fitting results from a meta-data analysis study using experimental findings from various clinical studies have estimated the highest viral concentration at 0.34 days after symptom onset ^14^.

Since data collected from both clinically confirmed COVID-19 cases and SARS-CoV-2 RNA abundance in wastewater are time series data, their relationships could be examined through time-series data analyses in particular cross-correlation. Removing the trend or seasonality of time-series data sets to achieve stationarity is an important prerequisite process to avoid spurious correlation ^15^. However, many of the wastewater surveillance studies that correlated SARS-CoV-2 in wastewater and COVID-19 cases in the community did not prewhiten the time series data ^16–19^. As a result, the strong correlation coefficients observed could be attributed to trend or seasonality instead of the actual correlation of variation between the two types of data sets.

In this study, we re-analyzed previously collected time-series data on daily wastewater SARS-CoV-2 RNA abundance and clinical COVID-19 cases in a large metropolitan area to demonstrate the importance of prewhitening when conducting cross-correlation analysis. Both SARS-CoV-2 RNA concentration and its normalized abundance were subjected to time-series cross-correlation analysis to determine any presence of lags between the corresponding daily clinical case numbers observed in the community. Also, normalization strategies were compared to distinguish the best improvement of correlation with the daily clinical case numbers.

## 2. Materials and Methods

### 2.1. Wastewater sampling, viral precipitation, RT process, and qPCR assays

Wastewater sampling, processing, and RT-qPCR quantification of SARS-CoV-2 RNA and several RNA viruses in wastewater were previously described in detail in Li *et al*. ^20^, which are briefly summarized in the following. The two largest wastewater treatment plants (WWTP), Sand Island (SI) and Honouliuli (HO) in the City and County of Honolulu, were selected to collect wastewater samples to represent the wastewater of the community. Untreated primary influent wastewater samples were collected by daily flow-adjusted composite sampling from the SI and HO WWTPs from August 27th, 2020 to October 4th, 2020 (i.e. Day 0 to 38, each WWTP with *n* = 39). All daily collected wastewater samples were thoroughly mixed and subsequently centrifuged to collect suspended solids and supernatant, which were referred to as solid and liquid fractions of the wastewater samples, respectively. The liquid fractions were first treated by the polyethylene glycol (PEG) precipitation method ^21^ to concentrate and pellet viral particles in the liquid fractions. The precipitated pellets from the liquid fractions as well as the solid fractions were subjected to viral RNA extraction. All extracted viral RNA samples were reverse transcribed with random hexamer N6, and the produced cDNA samples were analyzed by qPCR assays targeting SARS-CoV-2 (the N1 and N2 assays ^22^ and the E gene assay ^23^) and fecal RNA viral surrogates (F+ RNA coliphages Group II (G2) and Group III (G3) ^24^, and pepper mild mottle virus (PMMoV) ^25^).

### 2.2. Data analysis

All data analyses used both the log-transformed SARS-CoV-2 concentration data determined by the three qPCR assays (i.e. log N1, log N2, and log E) and its abundances normalized by the three fecal viral indicators (i.e. log (N1/G2), log (N2/G2), log (E/G2), log (N1/G3), log (N2/G3), log (E/G3), log (N1/PMMoV), log (N2/PMMoV) and log (E/PMMoV)). Daily new COVID-19 case counts for Honolulu were sourced from the local COVID-19 dashboard by the Hawaii Emergency Management Agency. The study only used publicly available data at the population level, and thus required no IRB review. Cross-correlation was used to examine the time-lagged association between new clinical COVID-19 cases in the community and wastewater SARS-CoV-2 abundance. A prewhitening process was applied to all time-series data, including COVID-19 clinical case numbers and the wastewater SARS-CoV-2 wastewater abundance, to remove trends. The wastewater SARS-CoV-2 abundance data were prewhitened by log transformation followed by the first difference, and the clinical case data were prewhitened by the first difference. The original and the prewhitened data were tested for normality by using Shapiro-Wilk test ^26^. Mann-Kendall test ^27, 28^ was used for the assessment of trend significance before and after the prewhitening to verify successful removal of trends.

Cross-correlation of original and prewhitened SARS-CoV-2 concentration and their normalized abundance in liquid or solid fractions and the daily new clinical COVID-19 cases were analyzed for the SI and HO WWTPs separately. The Cross-Correlation Functions (CCF) function in the R environment was used with a maximum lag of six days. A positive lag indicates that the SARS-CoV-2 concentration or normalized abundance was leading the clinical cases. Positive coefficients indicate a positive relationship between the SARS-CoV-2 concentration or normalized abundance and clinical cases.

Results of the cross-correlation analysis were visualized by heatmaps and boxplots. Additionally, correlation coefficients from the cross-correlation analysis were compared depending on normalization strategies by using *p*-values obtained from the pairwise *t*-test and were adjusted by the Benjamini and Hochberg correction ^29^ to determine which normalization strategy showed the best improvement. All statistical analyses and data visualization were conducted in R 4.2.1 ^30^ by using the packages *tidyverse* 1.3.1 ^31^, *ggpubr* 0.4.0 ^32^, *scales* 1.1.1 ^33^, and *rstatix* 0.7.0 ^34^.

## 3. Results

### 3.1. Importance of prewhitening on cross-correlation

Our previous study ^20^ detected downward trends for both daily clinical COVID-19 case numbers in the community and SARS-CoV-2 RNA abundances (both with or without normalization by fecal RNA viral markers) in the wastewater samples. The observed downward trends were the result of a public health lockdown implemented to counter the COVID-19 outbreak on the Island of Oahu. Therefore, the time-series data of raw wastewater SARS-CoV-2 RNA concentration and its normalized relative abundance, as well as the daily fluctuation of clinical case numbers all need to be prewhitened in order to be de-trended before cross-correlation analysis. The Shapiro-Wilk test showed that only 25 out of 48 of the original data (*p* = 0.08 ± 0.10) were normally distributed, but all of the prewhitened data (*p* = 0.39 ± 0.24) were normally distributed (Table S1). The Mann-Kendall test results confirmed the removal of trends from both SARS-CoV-2 abundance data from the wastewater and the daily clinical case data after the prewhitening (Table S1).

The prewhitened SI and HO WWTPs time-series SARS-CoV-2 concentration data were first compared with the prewhitened daily clinical case numbers, which showed only weak correlations (either positive or negative) (Figure 1). The only significant correlation was observed at a two-day lag of the prewhitened daily clinical case numbers (*x_t_+2*) behind the prewhitened daily wastewater SARS-CoV-2 concentrations of log N1 (*r* = 0.38, *p* = 0.019) in the liquid fraction of HO WWTP (Figure 1F). No significant correlations were found from any other lags from the two WWTPs with the prewhitened daily clinical case numbers, as indicated by the boxplots falling under the 95% confidence level (Figures 1BF).

Cross-correlation of the original non-stationary time-series SARS-CoV-2 data was also performed to illustrate the potential for spurious correlation (Figures 1CDGH). Both time-series concentration data (liquid and solid fractions) from SI (Figures 1CD) and HO (Figures 1GH) WWTPs showed all positive correlation coefficients and the majority of the correlation analyses showed statistically significant correlations with *p*-values less than 0.05 (SI: 21 out of 42 analyses; HO: 27 out of 42 analyses) with the original daily clinical case numbers (*x_t+h_*, *h* = lag number). Because the normality assumption of cross-correlation analysis was not met, these high positive correlation coefficients are considered spurious and false positive.

**Figure 1.**
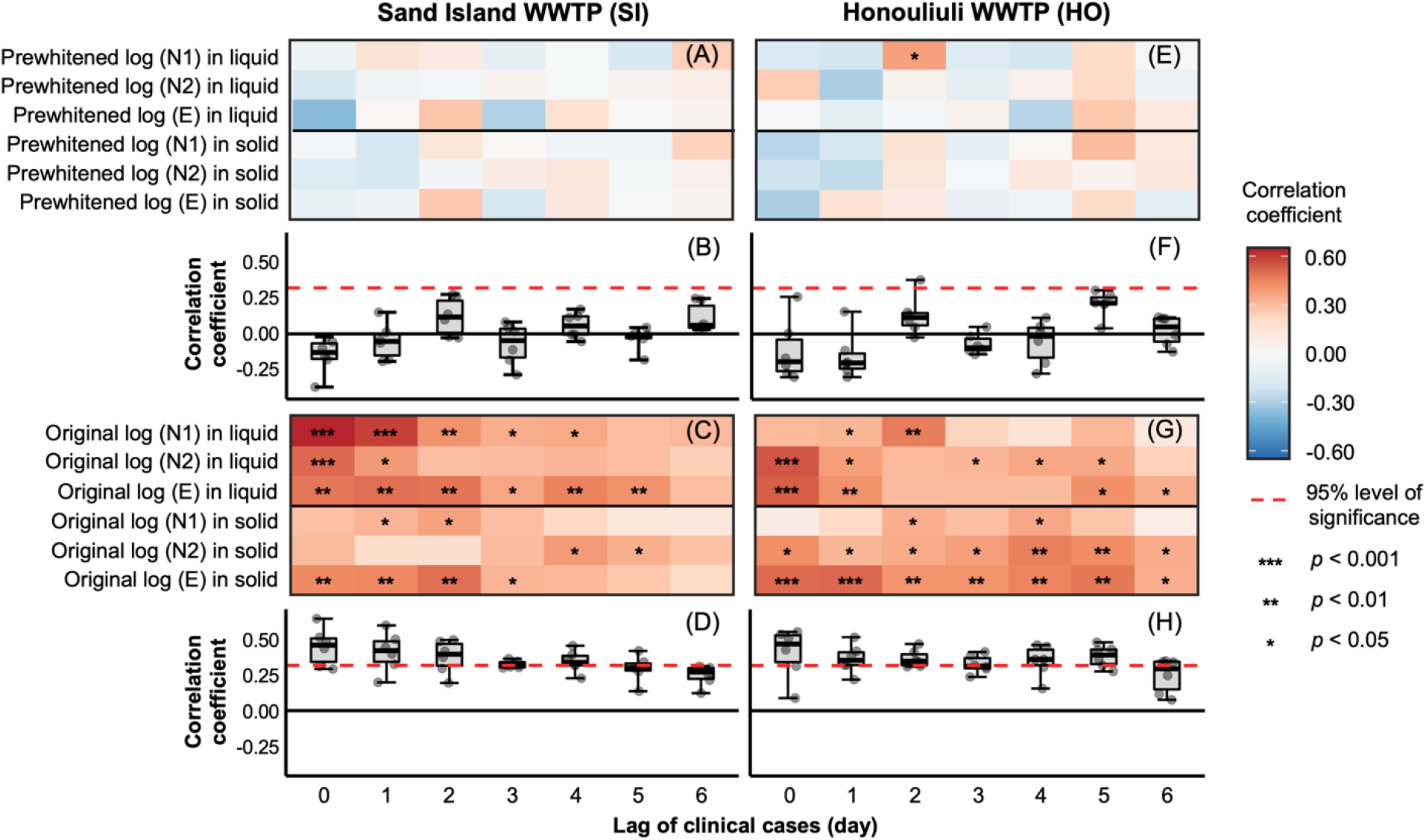
Cross-correlation between with and without prewhitening by the first difference of daily new clinical COVID-19 case numbers and measured SARS-CoV-2 RNA concentration (log 10 transformed) in wastewater samples from Sand Island (Non-prewhitened: A, B; prewhitened: C, D) and Honouliuli (Non-prewhitened: E, F; prewhitened: G, H). Red dashed lines represent a 95% level of significance and the *p*-value of the correlation less than 0.05 are displayed as asterisks. The middle, upper, and lower lines in the box of the boxplot represent the median, 25^th^, and 75^th^ percentiles, respectively, and the whiskers represent the largest and smallest values outside of the interquartile range.

### 3.2. Impact of normalized abundance on cross-correlation

The concentrations of SARS-CoV-2 RNA measured from the samples of the two WWTPs are expected to be impacted by various processes during wastewater sampling and sample processing, including total fecal discharge in the area, sewer collection in the WWTPs, wastewater viral precipitation method, and molecular quantification steps. The resulting variations could be potentially mitigated by normalizing data to various endogenous fecal viral indicators (e.g. log N1/G2) ^20^. The normalized daily wastewater SARS-CoV-2 RNA data were analyzed via cross-correlation with the prewhitened daily clinical case numbers. For wastewater samples from the SI WWTP, the normalization strategy produced significantly different cross-correlation patterns against different time lags (Figure 2) than those without normalization (Figure 1).

The most obvious improvement in the cross-correlation coefficient was observed at a two-day lag (*x*_t+2_, Figures 2BDF), with a range of *r* = −0.03-0.45 (0.23 ± 0.13). The average cross-correlation coefficients were increased from 0.12 to 0.33, 0.19, and 0.16 for G2, G3, and PMMoV normalizations, respectively, which showed an average of 0.11 ± 0.09 increase than those without normalization (Figures 2BDF). Among all combinations, the best correlation coefficient (*r* = 0.45, *p* = 0.004) was observed between the log (E/G2) in the liquid fraction and the daily clinical case numbers. The normalization strategy showed a higher improvement of correlation coefficients in liquid fractions (0.14 improved) than in the solid fractions (0.07 improved) compared to the raw data.

At the two-day lag, statistically significant correlations were observed more frequently with the normalized SARS-CoV-2 abundance data than with the raw data. For example, at the SI WWTP, both log (N1/G2) and log (E/G2) showed statistically significant correlation coefficients in the liquid (*r* = 0.38 (*p* = 0.017) and *r* = 0.45 (*p* = 0.004), respectively) and solid fractions (*r* = 0.32 (*p* = 0.048) and *r* = 0.38 (*p* = 0.018), respectively). In contrast, there was no statistically significant correlation between clinical cases and raw wastewater SARS-CoV-2 concentration data at SI WWTP (Figure 1A). G3 normalization showed two statistically significant correlations from log (N1/G3) and log (E/G3) (*r* = 0.33 (*p* = 0.043) and *r* = 0.33 (*p* = 0.044), respectively) in the liquid fractions. PMMoV normalization showed only one statistically significant correlation coefficient from the log (E/PMMoV) in the solid fraction (*r* = 0.33, *p* = 0.043).

**Figure 2.**
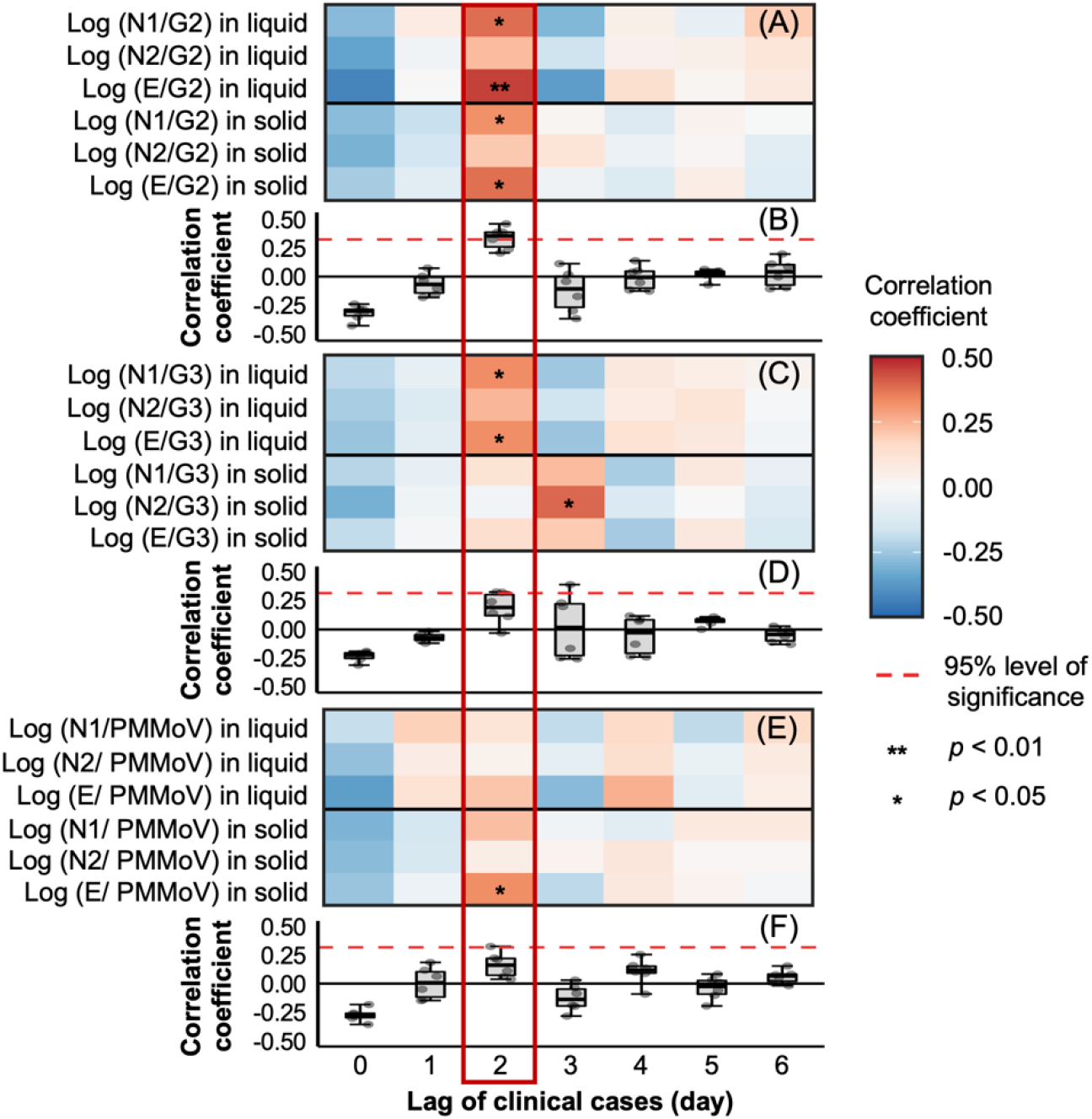
Cross-correlation between the prewhitened COVID-19 new case numbers and the prewhitened SARS-CoV-2 RNA normalized abundance in wastewater samples from the SI WWTP. The normalized abundance was calculated by dividing SARS-CoV-2 RNA abundance by F+ RNA coliphage Group II (A, B), Group III (C, D), and PMMoV (E, F). All normalized abundances were transformed into log forms. Red dashed lines represent a 95% level of significance and the *p*-value of the correlation less than 0.05 are displayed as asterisks. The middle, upper, and lower lines in the box of the boxplot represent the median, 25^th^, and 75^th^ percentiles, respectively, and the whiskers represent the largest and smallest values outside of the interquartile range.

For the HO WWTP, the largest correlation coefficients from the cross-correlation between the normalized SARS-CoV-2 abundance and the daily clinical case numbers were also observed at a two-day lag (*x_t+2_*, Figures 3BD), which is similar to the results of SI WWTP. The average cross-correlation coefficients were increased from 0.13 to 0.21 and 0.24 for G2 and G3 normalizations, respectively, which showed an average of 0.10 ± 0.02 increase than those without normalization (Figures 3BD). The best correlation coefficient among all three normalization methods was observed between the log (N1/G2) in the liquid fraction and the daily clinical case numbers (*r* = 0.35, *p* = 0.029). Similar to the SI WWTP, the normalization strategy showed a larger improvement of correlation coefficients in liquid fractions (0.07 improved) than in the solid fractions (0.03 improved) compared to the raw data.

At the two-day lag, all normalized forms of log N1 in the liquid fractions (log (N1/G2): *r* = 0.35, *p* = 0.029; log (N1/G3): *r* = 0.35, *p* = 0.030; log (N1/PMMoV): *r* = 0.33, *p* = 0.043) showed statistically significant correlations between the daily clinical cases and the normalized SARS-CoV-2 RNA abundance from HO WWTP (Figures 3ACE). Although no statistically significant correlation was observed from solid fractions from any normalization strategies (*r* = −0.05-0.27, 0.16 ± 0.11), all correlation coefficients observed from the solid fractions were increased after normalization by G2 and G3.

The correlation coefficients of all genes from both liquid and solid fractions of HO WWTP were decreased when they were normalized by PMMoV with average correlation coefficients decreased from 0.13 to 0.09 at the two-day lag (Figure 3E). Interestingly, PMMoV normalized SARS-CoV-2 RNA abundances in the liquid fraction showed large correlation coefficients (*r* = 0.32 ± 0.06) at a five-day lag of daily clinical case numbers (Figure 3F) showing a statistically significant correlation with log (E/PMMoV) (*r* = 0.39, *p* = 0.014).

The better correlation exhibited by the SI WWTP than the HO WWTP could be due to the SI WWTP treating a much larger fraction of the City’s wastewater (58%) than the HO WWTP (24%), and hence is expected to be more representative of the community disease burden.

**Figure 3.**
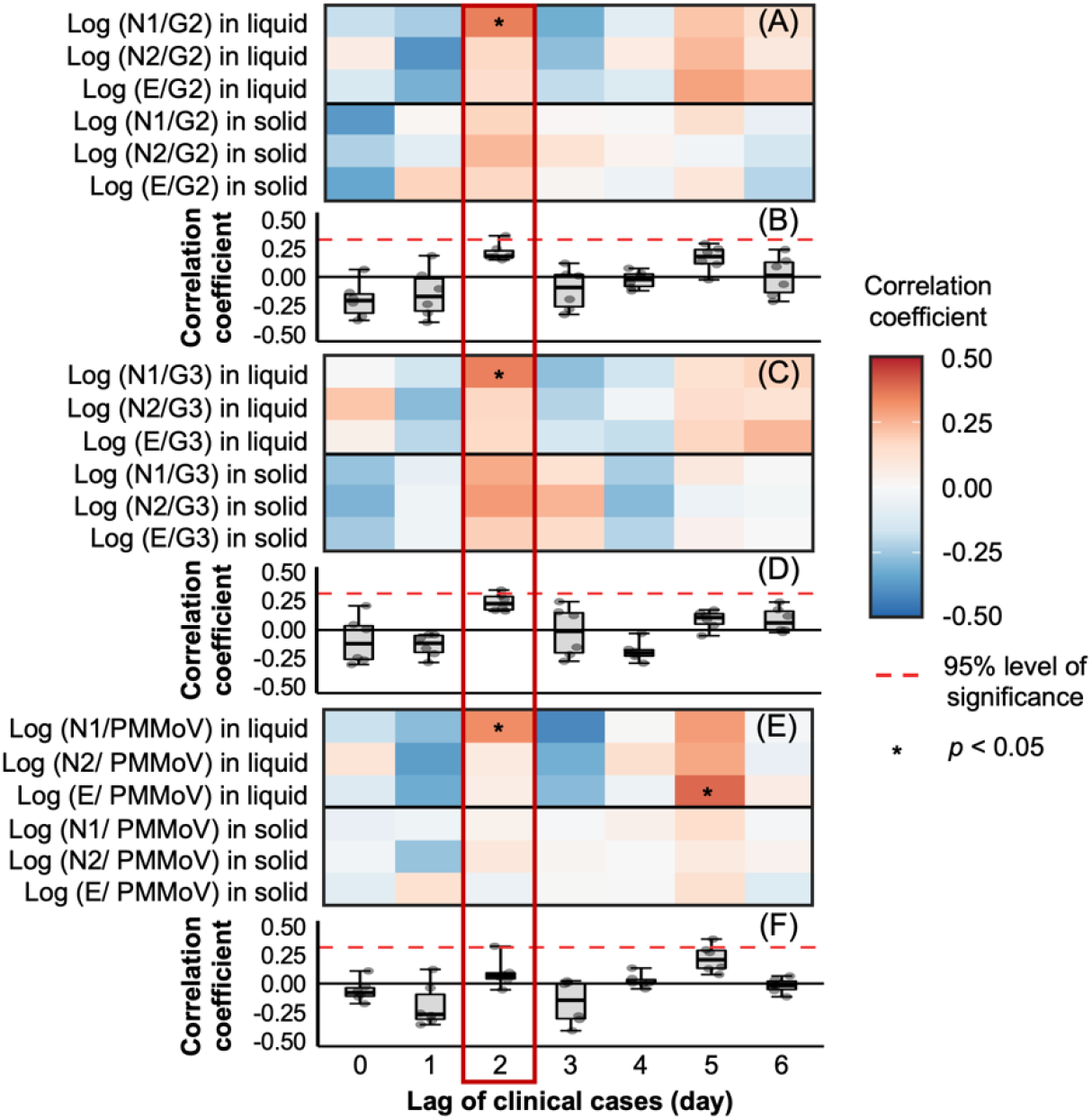
Cross-correlation between the prewhitened COVID-19 new case numbers and the prewhitened SARS-CoV-2 RNA normalized abundance in wastewater samples from the HO WWTP. The normalized abundance was calculated by dividing SARS-CoV-2 RNA abundance by F+ RNA coliphage Group II (A, B), Group III (C, D), and PMMoV (E, F). All normalized abundances were transformed into log forms. Red dashed lines represent a 95% level of significance and the *p*-value of the correlation less than 0.05 are displayed as asterisks. The middle, upper, and lower lines in the box of the boxplot represent the median, 25^th^, and 75^th^ percentiles, respectively, and the whiskers represent the largest and smallest values outside of the interquartile range.

### 3.3. Comparison of different normalization strategies for cross-correlation analysis

The correlation coefficients between the daily clinical case numbers and SARS-CoV-2 RNA abundance at a two-day lag of clinical cases were compared with respect to the normalization strategies by using a pairwise *t*-test (Figure 4). Among the three different endogenous fecal viral RNA controls used, only normalizing the data with G2 (*r* = 0.33 ± 0.10, *p* = 0.002) showed a statistically significant improvement of correlation coefficients in comparison to that using the raw data (*r* = 0.12 ± 0.13). While G3 (*r* = 0.19 ± 0.14, *p* = 0.396) and PMMoV (*r* = 0.16 ± 0.11, *p* = 0.198) mildly improved the correlation (Figure 4A), but not statistically. Furthermore, the G2 normalization showed a significantly larger correlation coefficient value than both G3 (*p* = 0.04) and PMMoV (*p* = 0.014) normalization methods.

**Figure 4.**
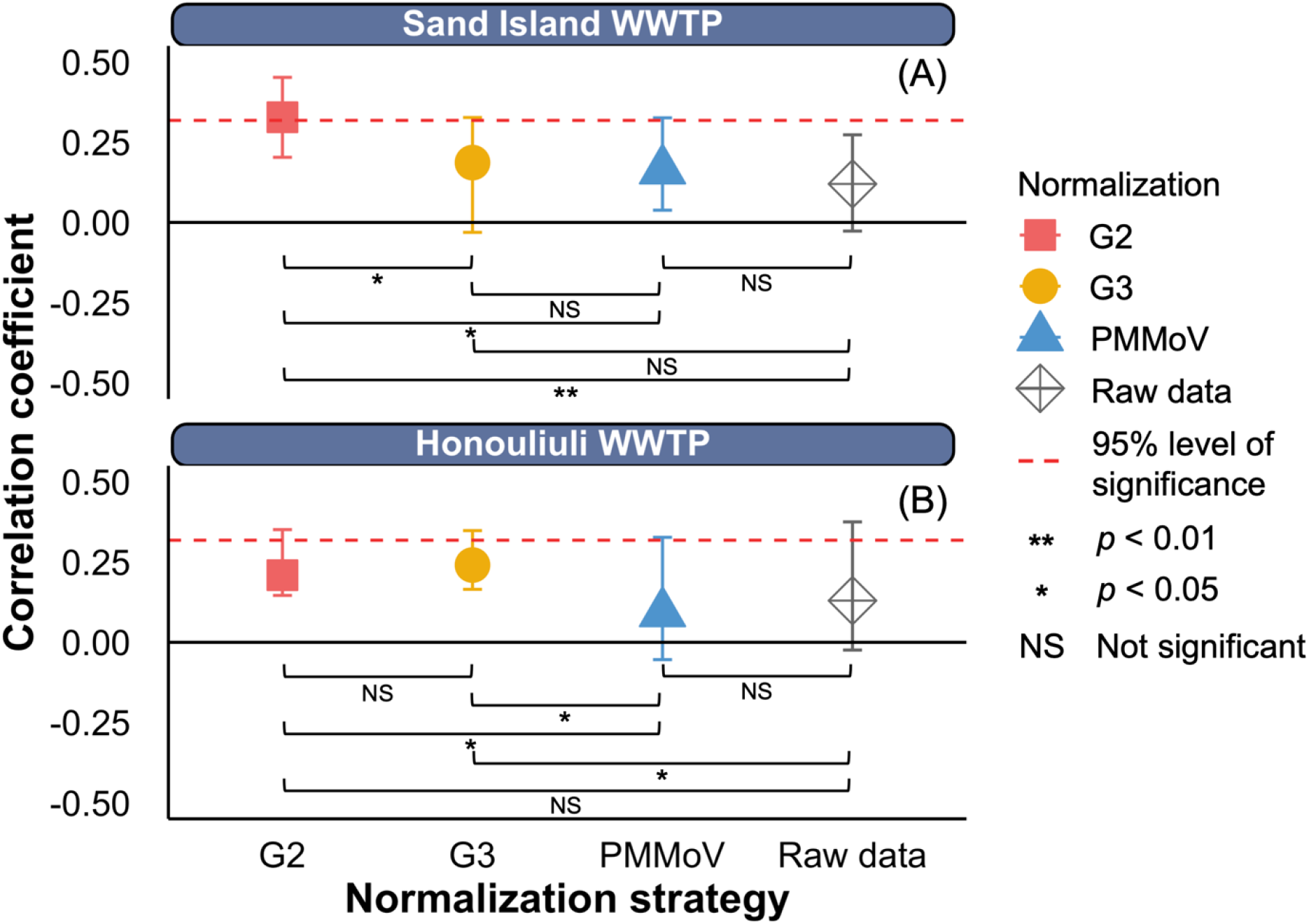
Cross-correlation between the first difference of daily new clinical COVID-19 case numbers in Honolulu County and normalized SARS-CoV-2 RNA concentration (log 10 transformed) by F+ RNA coliphage Group II, Group III, and PMMoV in wastewater samples from Sand Island (A) and Honouliuli (B) at a two-day lag of clinical cases. Red dashed lines represent a 95% level of significance and the whiskers represent the largest and the smallest values.

For the HO WWTP, the G3 (*r* = 0.24 ± 0.08, *p* = 0.032) normalization method was the only strategy that significantly improved the correlation coefficients (Figure 4B). Although G2 (*r* = 0.21 ± 0.08, *p* = 0.058) did not statistically improve the correlations with the raw data, it increased the mean values of correlation coefficients (*r* = 0.13 ± 0.14). However, PMMoV (*r* = 0.09 ± 0.13, *p* = 0.32) normalization even decreased the mean values of the correlation coefficients than the raw data. Moreover, G2 (*p* = 0.028) and G3 (*p* = 0.028) normalizations improved the correlation when compared with PMMoV normalization.

The overall results indicate that the normalization of SARS-CoV-2 RNA abundance improved the cross-correlations with the daily clinical case numbers. G2 normalization showed the largest improvement of cross-correlations in SI WWTP samples. On the contrary, the G3 normalization was the largest improvement of the cross-correlations in HO WWTP samples. When considering the liquid fractions only, the G2 normalization of log N1 from SI (*r* = 0.38, *p* = 0.017) and HO (*r* = 0.35, *p* = 0.029) WWTPs showed both significant correlations with the daily clinical case numbers.

## 4. Discussion

In our previous study ^20^, we observed simultaneous downward trends between SARS-CoV-2 RNA abundances (both with and without normalization by fecal viral markers) in wastewater samples from the SI and HO WWTPs and the daily clinical COVID-19 case numbers during a COViD-19 public health lockdown. This is congruent with previous observations where increases in wastewater SARS-CoV-2 RNA concentration corresponded with rapidly expanding COVID-19 outbreaks ^32–36^. The fine-scale temporal dynamics afforded by daily sampling also detected significant intra-day fluctuation of the wastewater SARS-CoV-2 RNA abundance, even within the same weeks. Many factors could have contributed to the observed intra-day fluctuation, including errors in wastewater sampling and sample analysis, variations in viral shedding by infected individuals, and daily fluctuations in disease burden in the community. Since similar trends were detected in the two replicate WWTPs, over multiple weeks, and regardless of normalization strategies, the former two (i.e. sampling and analysis errors and variation in viral shedding) are unlikely to entirely explain the observations.

Cross-correlation analysis of the time-series data of wastewater SARS-CoV-2 abundance and clinical case numbers could be used to infer potential association and determine if daily fluctuations in disease burden in the community contributed to the observed intra-day fluctuation. Many wastewater surveillance studies have compared wastewater SARS-CoV-2 abundance with clinical case data in the community through correlation analysis ^14–17^. However, few previous studies have conducted pre-whitening treatment to achieve stationarity of the time-series data. Stationarity in time-series data indicates consistency of the distribution (mean and variance) over time ^45^, and non-stationary time-series data can often lead to spurious outcomes in correlation analyses. This was clearly demonstrated when we observed significant spurious cross-correlations with the original data that contain trends (Figure 1 CDGH). Similar phenomenon may explain reports of wastewater surveillance showing strong correlations to clinical cases, especially when the studies were conducted during a period when COVID-19 clinical cases were continuously increasing or decreasing ^32, 46, 47^. Therefore, prewhitening the data for wastewater surveillance to meet the stationarity requirement of cross-correlation analysis must be practiced in order to identify the actual association between data sets.

After prewhitening the data, the cross-correlation coefficients decreased significantly, and the overall patterns with respect to the time lag also changed drastically (Figure 1). For example, at zero-time lag, cross-correlation using the original data detected the best positive correlation, whereas the prewhitened data actually detected some negative correlations. The low levels of correlation detected are not entirely unexpected, considering the extraordinary complexity involved in collecting the wastewater SARS-CoV-2 RNA data and resulting variations. Many factors, including varying fecal discharge by infected individuals, dilution and fluctuation during transportation in sanitary sewers, and wastewater sample collection and processing, could have contributed to the variations in SARS-CoV-2 RNA in wastewater samples. The molecular quantification processes could also introduce additional variations to the results; for example, RNA recovery during sample extraction could have varying efficiencies, and subsequent reverse transcription and qPCR quantification could introduce additional biases.

The incorporation of normalization strategies and the resulting relative abundance of SARS-CoV-2 RNA in the wastewater samples led to the identification of a two-day lag showing the best correlation (Figures 2 and 3). Given the complex and multi-step process required for quantifying SARS-CoV-2 RNA in wastewater, the selection of endogenous viral RNA control for global normalization is also important. Amongst the three fecal RNA viruses as endogenous controls tested in this study, normalization by G2 provided the most significant improvement in correlation between wastewater SARS-CoV-2 RNA abundance and clinical new cases followed by G3. Cole *et al*. found that G2 had the highest proportion among the total F+ RNA groups in WWTP samples (51.9%) and G2 was found more in human-impacted wastes than G3 ^48^. This supports our results of G2 normalization of SARS-CoV-2 RNA abundance having higher correlation coefficients compared to G3. On the other hand, PMMoV only provided marginal improvement in correlation. This difference could be attributed to their respective sources in human feces where G2 and G3 are inherently linked with fecal coliforms while PMMoV is subjected to dietary variation in pepper consumption. Some previous wastewater surveillance studies that used PMMoV for the SARS-CoV-2 abundance normalization also reported that the PMMoV did not improve the correlation with the clinical cases ^49-51^.

The significant cross-correlation between the normalized abundance of daily wastewater SARS-CoV-2 and new clinical cases in the community is highly intriguing. Since the COVID-19 pandemic, weekly intra-day oscillations in new clinical case numbers have been widely observed in communities across the globe ^37^. One school of thought is that these weekly intra-day oscillations are primarily a reflection of diagnostic and reporting biases ^38, 39^, while a competing theory is that this could be caused by actual disease transmission dynamics caused by weekly behavior patterns ^40-42^. The strong correlation between the intra-day fluctuation and weekly oscillation of wastewater SARS-CoV-2 RNA abundance and clinical case numbers that we observed in this study suggest that the observed weekly oscillation of clinical cases may be indeed a true reflection of disease burden in the community, in addition to contributions from clinical sampling and reporting biases and errors.

Since the average turnaround for clinical testing during the study period was approximately one day, with the assumption of one day lag between symptom onset and specimen collection, the observed two-day lag in cross-correlation analysis indicates that the wastewater SARS-CoV-2 viral RNA abundance may be synchronizing with symptom development of new COVID-19 cases in the community. Studies at the early stage of the pandemic, which likely experienced clinical testing delays, have reported the detection of the SARS-CoV-2 viral RNA in wastewater about one week ahead of reported clinical cases in the communities ^33, 36^, while a study reported wastewater sludge SARS-CoV-2 concentration leading the specimen collection by 0-2 days ^22^. The apparent synchronous correspondence supports the possibility of using wastewater for early detection of viral transmission in communities, as viral shedding can start 3-5 days before and peaks around symptom onset ^43^ and asymptomatic infections and often lead to symptomatic infections and represent a significant portion of the overall community disease burden ^44^.

In this study, both the solid and liquid fractions of the same wastewater samples were analyzed separately, and SARS-CoV-2 normalized abundance data in both fractions showed similar cross-correlation patterns (Figures 2 and 3). In the previous study ^18^, the solid fractions contained higher per mass concentration of SARS-CoV-2 RNA than the liquid fractions, while the normalized abundances between the two fractions were quite similar. The normalized abundance of SARS-CoV-2 RNA in liquid fractions exhibited slightly stronger correlations with the clinical COVID-19 case numbers in the community than the normalized abundance of SARS-CoV-2 in the solid fraction. This could be attributed to the more complex matrix effects in the solid fractions than the liquid fractions, as indicated in our previous study where lower recovery and higher variation of the spiked bovine coronavirus (BCoV) as exogenous control were observed in the solid fractions than in the liquid fractions ^18^.

It is important to note that all three quantification assays showed similar cross-correlation patterns between normalized SARS-CoV-2 RNA abundance in wastewater and clinical case numbers in the community. While the SARS-CoV-2 RNA genome contains a single copy of N and E genes, our previously published study ^18^ and many other studies ^33, 56^ have shown that different assays usually generate different abundance levels, indicating that the molecular quantification processes have variations. Nevertheless, all three assays were able to reveal strong correlations at a two-day lag between normalized SARS-CoV-2 RNA abundance in the wastewater and community disease burden, with the N1 gene assay providing the highest correlation coefficients in both tested WWTPs (Figures 2 and 3). Even though the E gene is considered the least specific PCR target for SARS-CoV-2 detection due to homologous sequence similarities with other coronaviruses together with recurrent mutations ^57^, strong positive cross-correlation patterns were also determined at a two-day lag from both SI and HO WWTPs which were similar to those of N1 and N2 genes.

## 5. Conclusions

This study demonstrated the importance of prewhitening to remove the trends of the daily fluctuation of wastewater surveillance data and the clinical case numbers before cross-correlation analysis of the time-series data sets to avoid spurious correlations. We also observed that normalization strategies to account for various variations in the process are helpful in improving cross-correlation analysis. Amongst the various normalization strategies, SARS-CoV-2 RNA abundances normalized with F+RNA coliphage G2 provided the best correlation coefficients in this study. This enabled us to show that the daily clinical case numbers were two days behind the SARS-CoV-2 RNA detection in wastewater, which supports the notion that wastewater surveillance has the potential to provide earlier detection of SARS-CoV-2 signal than clinical diagnosis. Most interestingly, the strong cross-correlation between the intra-day fluctuation and weekly oscillation of wastewater SARS-CoV-2 RNA abundance and clinical cases suggest that the observed weekly oscillation of clinical cases is likely caused (at least partially) by disease burden fluctuation in the community in additional to clinical sampling and reporting biases and errors, which requires further research to delineate their respective contributions.

## Acknowledgments

This material is based upon work supported by the National Science Foundation under Grant No. CBET-2027059.

