## Supplemental Table S1, Figure S1, Figure S2 for "Prewhitening and Normalization Help Detect a Strong Cross-Correlation Between Daily Wastewater SARS-CoV-2 RNA Abundance and COVID-19 Cases in a Community"

### Supplementary data

**Table S1**. Slope, trend, and normality test results of before and after prewhitening of SARS-CoV-2 abundance (both raw and normalized) from wastewater and daily clinical case numbers

| Sample | Analysis | Prewhitening | Average | Standard Deviation | Maximum value | Minimum value |
|---|---|---|---|---|---|---|
| Honouliuli WWTP ($n$ = 24) | Slope | Before | -0.056 | 0.024 | -0.021 | -0.100 |
| | | After | 0.002 | 0.005 | 0.013 | -0.010 |
| | Mann-Kendall trend test $p$-value[a] | Before | 0.013 | 0.034 | 0.137 | 0.000 |
| | | After | 0.809 | 0.159 | 1.000 | 0.435 |
| | Shapiro-Wilk test $p$-value[b] | Before | 0.101 | 0.104 | 0.445 | 0.002 |
| | | After | 0.443 | 0.252 | 0.963 | 0.093 |
| Sand Island WWTP ($n$ = 24) | Slope | Before | -0.055 | 0.020 | -0.014 | -0.100 |
| | | After | 0.003 | 0.005 | 0.011 | -0.007 |
| | Mann-Kendall trend test $p$-value | Before | 0.000 | 0.001 | 0.007 | 0.000 |
| | | After | 0.799 | 0.169 | 1.000 | 0.443 |
| | Shapiro-Wilk test $p$-value | Before | 0.069 | 0.100 | 0.449 | 0.002 |
| | | After | 0.344 | 0.236 | 0.831 | 0.055 |
| Daily clinical case numbers ($n$ = 1) | Slope | Before | - | - | -3.5081 | |
| | | After | - | - | 0.2655 | |
| | Mann-Kendall trend test $p$-value | Before | - | - | 0.000 | |
| | | After | - | - | 0.308 | |
| | Shapiro-Wilk test $p$-value | Before | - | - | 0.002 | |
| | | After | - | - | 0.436 | |

[a] The null hypothesis of Mann-Kendall trend test is that the data has no trend or serial correlation structure throughout the observed time points.

[b] The null hypothesis of Shapiro-Wilk test is that the data is normally distributed.

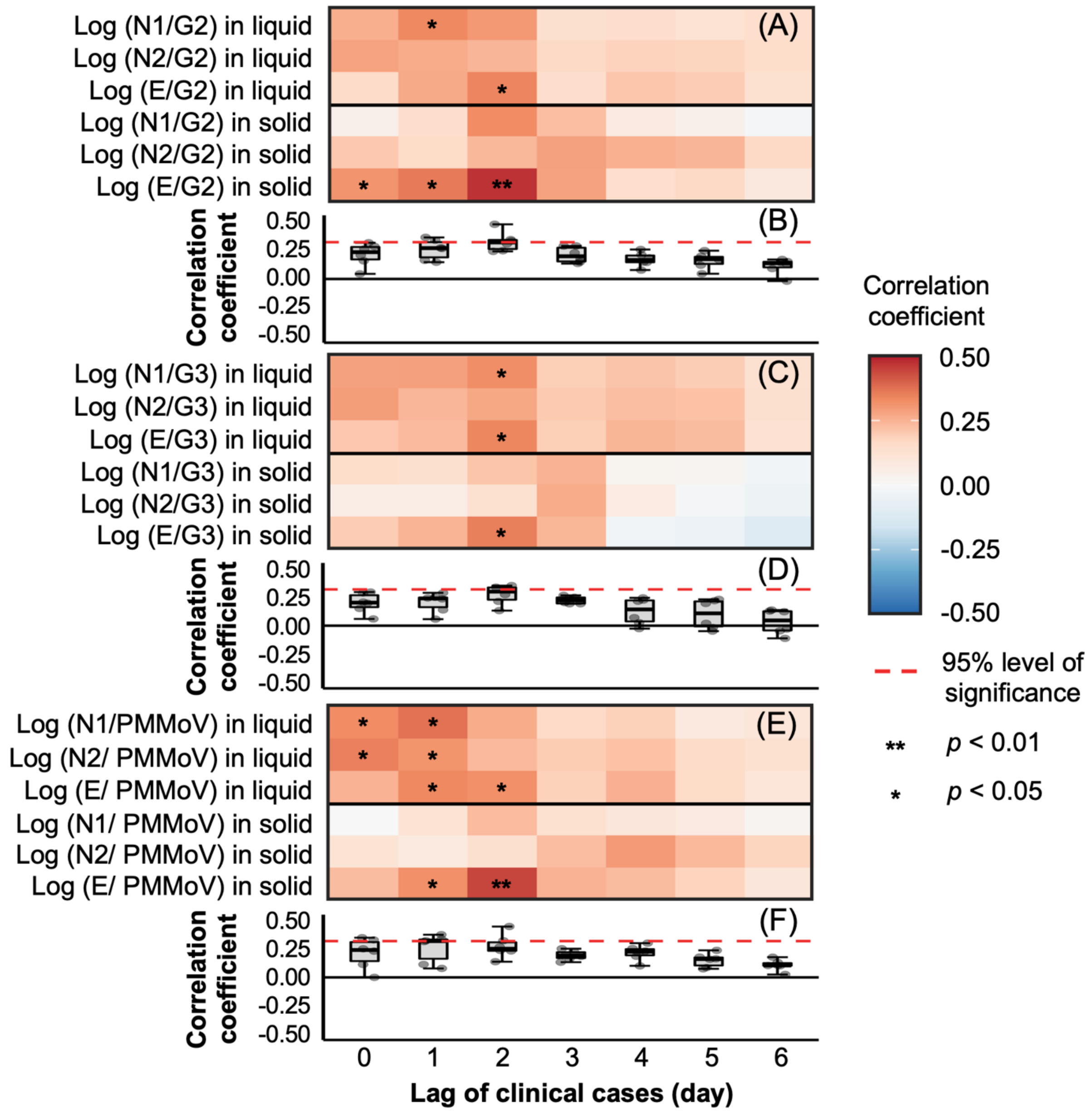


**Figure S1.** Cross-correlation between the non-prewhitened COVID-19 new case numbers and the non-prewhitened SARS-CoV-2 RNA normalized abundance in wastewater samples from the SI WWTP. The normalized abundance was calculated by dividing SARS-CoV-2 RNA abundance by F+ RNA coliphage Group II (A, B), Group III (C, D), and PMMoV (E, F). All normalized abundances were transformed into log forms. Red dashed lines represent a 95% level of significance and the *p*-value of the correlation less than 0.05 are displayed as asterisks. The middle, upper, and lower lines in the box of the boxplot represent the median, 25$^{th}$, and 75$^{th}$ percentiles, respectively, and the whiskers represent the largest and smallest values outside of the interquartile range.

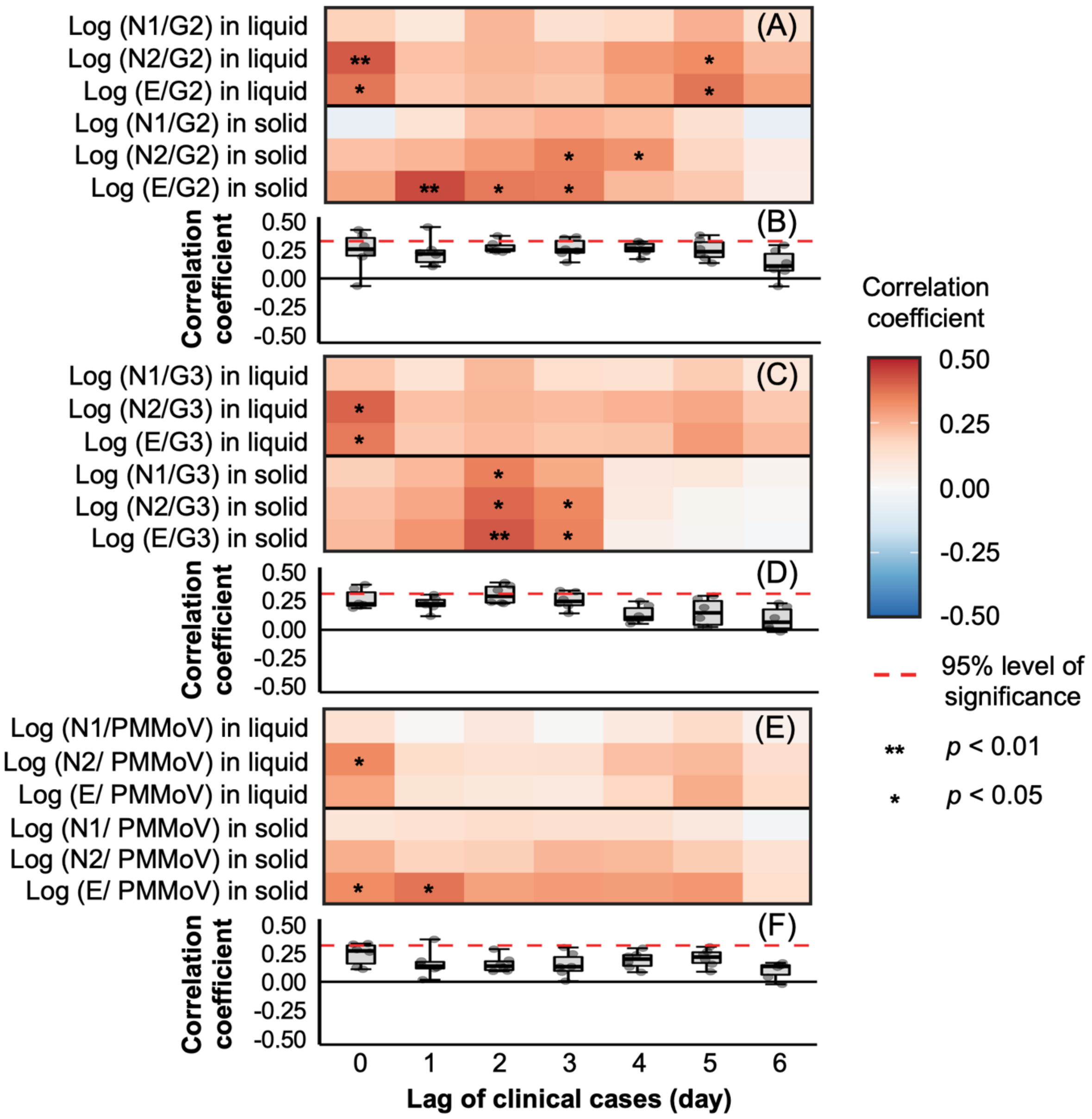


**Figure S2.** Cross-correlation between the non-prewhitened COVID-19 new case numbers and the non-prewhitened SARS-CoV-2 RNA normalized abundance in wastewater samples from the HO WWTP. The normalized abundance was calculated by dividing SARS-CoV-2 RNA abundance by F+ RNA coliphage Group II (A, B), Group III (C, D), and PMMoV (E, F). All normalized abundances were transformed into log forms. Red dashed lines represent a 95% level of significance and the *p*-value of the correlation less than 0.05 are displayed as asterisks. The middle, upper, and lower lines in the box of the boxplot represent the median, 25th, and 75th percentiles, respectively, and the whiskers represent the largest and smallest values outside of the interquartile range.
